# Comparative Proteomics Reveals Cell-Type-Specific Heterogeneity of Human Neural Cell-Derived Extracellular Vesicles

**DOI:** 10.64898/2026.10.29.762880

**Authors:** Xinqian Chen, Chih Hung Lo, Jialiu Zeng

## Abstract

Extracellular vesicles (EVs) are increasingly explored as biomarkers and as drug delivery vehicles for the brain, yet how EV cargo differs among neural cell types at baseline remains poorly defined. This gap limits both the interpretation of disease-associated EV changes and the rational choice of EV source for therapeutic use. Here, we compared EVs from human neuronal-like SH-SY5Y cells (N-EV), astrocytes (AS-EV), and HMC3 microglia (MG-EV) side by side using tandem mass tag (TMT)-based quantitative proteomics, with all samples analyzed in a single multiplexed batch. EV identity was supported by TEM, nanoparticle tracking analysis, and canonical MISEV2023 markers. Size, zeta potential, and protein content per vesicle were comparable across EV types, whereas N-EV showed the highest particle concentration and protein yield. Enrichment analyses and consensus clustering showed cargo consistent with parental-cell functions: neuronal connectivity and structural remodeling in N-EV, extracellular support and cellular interaction in AS-EV, and immune surveillance in MG-EV. Cross-comparison also revealed patterns beyond these canonical roles, including complement-associated cargo shared by N-EV and MG-EV and a Glial-EV signature of chromatin-associated and epigenetic regulatory proteins shared by AS-EV and MG-EV. We further defined cell-type-specific signature proteins; interaction networks of the larger sets were consistent with the enrichment results. Overall, these data provide a baseline comparative reference of EV proteomes from human neural cell models for interpreting disease-associated alterations, selecting EV sources for therapeutic use, and generating hypotheses on selective cargo sorting.

## Introduction

The central nervous system (CNS) primarily comprises specialized cell types including neurons, astrocytes, and microglia, which maintain brain homeostasis through continuous intercellular communication. Extracellular vesicles (EVs) have emerged as key mediators of such communication, delivering bioactive cargo including proteins, lipids, nucleic acids, and metabolites between cells under both physiological and pathological conditions [1]. Growing evidence indicates that EVs participate in diverse biological processes, including intercellular signaling, immune regulation, brain homeostasis, and cellular responses to injury, and are increasingly recognized as biomarkers and therapeutic delivery vehicles because of their ability to cross the blood-brain barrier [2–5]. Importantly, EV cargo reflects the identity and physiological state of its cells of origin, making EV proteomics a powerful approach for investigating cell-specific biology and identifying disease-associated molecular alterations [5]. To date, proteomic studies of CNS cell-derived EVs have focused predominantly on disease or activation states. Proteomic profiling of brain-derived EVs from Alzheimer’s disease (AD) and control human brain tissue has revealed that AD EVs are enriched with pathogenic proteins, including amyloid-β, tau, apolipoprotein E, and α-synuclein, and machine learning identified a panel of EV proteins that distinguished AD from control cases with 88% accuracy [6]. More recently, comparative proteomic analysis demonstrated that disease-associated proteomic alterations are more pronounced in brain-derived EVs than in the corresponding brain tissue, with elevated tau- and oxidative stress-related proteins and altered neuron-, astrocyte-, microglia-, and endothelial-associated EV surface markers, highlighting extensive remodeling of EV cargo during neurodegeneration [7].

Although physiological EVs have been included as control groups in disease-oriented studies, their baseline protein diversity across CNS cell types remains incompletely characterized. Differences in cellular protein composition and selective EV cargo sorting may contribute to this heterogeneity among EVs released by neurons, astrocytes, and microglia. Selective EV cargo loading is now recognized as a regulated process that can enrich or exclude specific protein classes rather than simply reflecting cellular abundance [8–10]. Comparative proteomic studies further demonstrate that EVs can display protein profiles distinct from their parental cells, with selective enrichment and depletion of specific proteins observed across multiple experimental systems, including neural stem cell-derived EVs [11–13]. Beyond simply reflecting the functions of their parent cells, such selective cargo sorting may enrich EVs with proteins associated with biological processes that are not readily predicted from the established roles of their cells of origin. Characterizing this heterogeneity may therefore identify shared and cell-type-associated EV protein signatures and provide insight into the potential roles of different CNS cell-derived EVs in cell-to-cell communication. Establishing such a baseline reference would help interpret disease-associated changes in EV protein composition. Recent studies have begun to characterize baseline CNS EV proteomes using mouse brain tissue [14] and induced pluripotent stem cell (iPSC)-derived neural cells [15], supporting source-dependent heterogeneity in EV cargo profiles. Understanding EV heterogeneity can also help guide therapeutic development and the use of EVs as drug delivery vehicles [16–18]. Astrocyte-derived EVs have been shown to enhance neuronal survival and electrophysiological function in vitro [19], while EVs from different cellular sources differ in properties relevant to brain delivery [20]. However, the proteins contributing to these effects and differences have yet to be systematically characterized. Comparative proteomic analysis of CNS cell-derived EVs may therefore identify candidate proteins associated with neuroprotective activity, cellular targeting, and uptake, thereby supporting the selection or engineering of EV sources for therapeutic applications.

In this study, we used tandem mass tag (TMT)-based multiplexed quantitative proteomics to characterize and compare the EV proteomes of three widely used human CNS cells, including SH-SY5Y human neuroblastoma cells, hAs human astrocytes, and HMC3 human microglia under baseline culture conditions. These cell models were selected to represent neurons, astrocytes, and microglia, three major CNS cell types involved in neuronal signaling, homeostatic support, and immune surveillance. By comparing their EV proteomes, we aimed to understand the shared and distinct molecular features of CNS cell-derived EVs under physiological conditions, explore how EV heterogeneity relates to their biological functions and potential therapeutic or drug delivery applications, and establish a baseline proteomic reference for future comparison with disease conditions.

## Materials and Methods

### Cell Culture and EV Isolation from Cell Media

SH-SY5Y human neuroblastoma cells (Cytion, #300154), hAs human astrocytes (Innoprot, P10251-IM), and HMC3 human microglia cells (Cytion, #300102) were cultured in Eagle’s Minimum Essential Medium/Ham’s F-12 (EMEM/F-12, 1:1), Human Astrocyte Medium (ScienCell, #1801), and EMEM (ATCC, #30-2003), respectively. All media were supplemented with 10% fetal bovine serum (FBS) and 1% penicillin/streptomycin (P/S), and vacuum-filtered through sterile 0.22 µm filters prior to use. Cells were maintained at 37 °C in a humidified 5% CO₂ incubator and passaged at 80–90% confluence.

To eliminate exogenous vesicular contamination from serum, EV-depleted FBS was prepared prior to conditioned medium collection. Briefly, FBS was ultracentrifuged at 100,000 × g for 18 h at4 °C in an Optima L-100XP ultracentrifuge (Beckman Coulter, Brea, CA, USA), and the supernatant was collected. For conditioned medium collection, cells were washed twice with phosphate-buffered saline (PBS), and the culture medium was replaced with the respective medium supplemented with EV-depleted FBS. Conditioned medium was collected after a minimum of 48 h of incubation and processed for EV isolation.

EVs were isolated from conditioned medium by differential ultracentrifugation. Conditioned medium was centrifuged at 2,000 × g for 20 min to remove cells, followed by 10,000 × g for 30 min to remove dead cells and large debris. The supernatant was filtered through a 0.22 µm syringe filter, and EVs were pelleted by ultracentrifugation at 100,000 × g for 90 min at 4 °C. The pellet was washed in PBS and re-pelleted at 100,000 × g for 90 min. The final EV pellet was resuspended in PBS and stored at −80 °C until further use.

### EV Characterization via Nanoparticle Tracking Analysis (NTA), Transmission Electron Microscopy (TEM), and Bicinchoninic Acid (BCA) Assay

The size distribution, particle concentration, and zeta potential of isolated EVs were determined by nanoparticle tracking analysis (NTA) on a ZetaView instrument (Particle Metrix Inc., Holly Springs, NC, USA). A 1 µL aliquot of each EV sample was diluted in double-distilled water (ddH₂O) to achieve 100–200 particles per frame, corresponding to an approximate dilution of 1:50,000–1:100,000.

Carbon-coated 200-mesh copper transmission electron microscopy (TEM) grids were glow-discharged at 15 mA for 30 s before sample application. A 4 µL aliquot of EV suspension was applied to each grid and allowed to adsorb for 5 min. Excess sample was wicked away with filter paper, and the grids were fixed in 2% paraformaldehyde (PFA) for 5 min. Grids were then washed twice on 50 µL drops of distilled water for 2 min per wash, with excess liquid wicked away between steps. Negative staining was performed using 1% uranyl acetate for 1 min. Excess stain was removed by wicking, and the grids were allowed to dry completely on filter paper before being transferred to a grid storage box. Imaging was performed using a JEOL JEM-1400 TEM operated at 80 kV. Digital micrographs were acquired using a Gatan Orius SC-1000 CCD camera.

EV protein concentration was determined using the bicinchoninic acid (BCA) protein assay. Briefly, EVs were lysed by mixing 5 µL of EV sample with 5 µL of RIPA buffer (Thermo Scientific, #89901) supplemented with phosphatase and protease inhibitors (Thermo Scientific, #78440) (100:1, v/v), followed by ultrasonication using a Sonic Dismembrator Model 100 (Fisher Scientific) for 3 s per cycle, for a total of 3 cycles. The lysate was then applied to a microplate for the BCA assay according to the manufacturer’s instructions.

### Proteomics Sample Preparation, TMT Labeling, and Fractionation

Biological replicates (n=2 per group) were prepared for multiplexed quantitative mass spectrometry. Proteins were buffer-exchanged using a 3 kDa molecular weight cutoff filter (Amicon Ultracel-3K; MilliporeSigma, Burlington, MA, USA) into 50 mM triethylammonium bicarbonate (TEAB, pH 8.0), and 10 µg of protein per sample was digested with an EasyPep Mini MS Sample Prep Kit (Thermo Fisher Scientific, #A40006) according to the manufacturer’s instructions, including reduction, alkylation, and overnight trypsin/Lys-C digestion at 37 °C. TMT 10-plex reagents (Thermo Fisher Scientific, #90110) were reconstituted in acetonitrile (ACN) and added to each digested sample. After 60 min, the reaction was quenched with 20% formic acid and 5% hydroxylamine, followed by solid-phase extraction cleanup and SpeedVac drying. Labeled samples were reconstituted in 3% ACN/0.2% trifluoroacetic acid (TFA), pooled, and fractionated using a Pierce High pH Reversed-Phase Peptide Fractionation Kit (Thermo Fisher Scientific, #84868) with step elution from 5–50% ACN in 0.1% triethylamine (TEA), yielding eight fractions for downstream analysis.

### LC-MS/MS Analysis, Database Searching, and Quantification

Dried fractions were reconstituted in 3% ACN/0.5% formic acid and injected onto a nano-LC column (75 µm × 28 cm, packed with 2.4 µm C18AQ particles) maintained at 50 °C. Peptides were separated over a 135 min gradient and analyzed on an Orbitrap Fusion Lumos Tribrid mass spectrometer (Thermo Fisher Scientific) operating in data-dependent top-speed mode (cycle time, 3 s). MS1 scans were acquired at a resolving power of 120,000 over m/z 375–1,500, and MS2 scans were acquired at a resolving power of 50,000 following higher-energy collisional dissociation (HCD) with a normalized collision energy of 37%. Raw data were searched with the SEQUEST HT algorithm within Proteome Discoverer (version 2.4, Thermo Fisher Scientific) against the Homo sapiens UniProt reference proteome (20,392 sequences) supplemented with a common contaminants database (cRAP). Search parameters included fully tryptic specificity with up to two missed cleavages, a precursor mass tolerance of 10 ppm, and a fragment mass tolerance of 0.02 Da. Carbamidomethylation of cysteine (+57.0215 Da) and TMT 10-plex labeling (+229.1629 Da) at lysine residues and peptide N-termini were set as static modifications, and oxidation of methionine (+15.9949 Da) was set as a dynamic modification. Peptide-spectrum matches were validated with Percolator and filtered to a 1% false discovery rate (FDR) at both the peptide and protein levels. Protein quantification was performed using TMT reporter ion intensities at the MS2 level, with a co-isolation threshold of 50% and an average reporter S/N threshold of 10, with abundances normalized to total peptide amount across channels.

### Differential Expression and Proteomic Analysis

Proteins were filtered based on the following criteria: (1) protein FDR confidence set to “High”; (2) experiment-level q-value < 0.01; (3) contaminant proteins were removed; (4) proteins identified by only one peptide were excluded; and (5) proteins lacking abundance values were removed. The filtered protein abundance values were then used for downstream differential expression analysis. Differential expression analysis was performed using the limma package [21,22] in R. A linear model was fitted to log₂-transformed protein intensities with empirical Bayes moderation. An omnibus F-test was first applied across all three groups to identify common differentially expressed proteins (DEPs), defined as proteins with an adjusted p-value < 0.01 (Benjamini-Hochberg correction) and a |log₂ fold change (FC)| ≥ 1. Subsequently, pairwise comparisons (AS-EV vs N-EV, AS-EV vs MG-EV, and MG-EV vs N-EV) were performed using limma contrasts to identify group-specific DEPs. Principal component analysis (PCA) was performed to assess sample clustering and visualize expression profiles. Volcano plots were generated to visualize DEPs between AS-EV, N-EV, and MG-EV groups.

### Subcellular Compartment Classification

Each protein was annotated with subcellular localization information retrieved from the UniProt database. Unclassified proteins were annotated with Gene Ontology (GO) Cellular Component (GO:CC) terms or manually annotated if no term could be assigned. Proteins were assigned to functional compartment classes based on their annotated localization (plasma membrane, endosome, lysosome, endoplasmic reticulum, Golgi apparatus, mitochondrion, cytoskeleton, secreted, cytoplasm, and nucleus). Proteins annotated to three or more compartments, or to two compartments not covered by the predefined combinations, were classified as “other multiple compartments.” The number of up- and downregulated DEPs within each functional compartment class was visualized using the ggplot2 package of R.

### Functional Enrichment and Pathway Analysis

GO enrichment analysis for Biological Process (BP) and Cellular Component (CC) terms, and Kyoto Encyclopedia of Genes and Genomes (KEGG) pathway analysis were performed on DEPs from each pairwise comparison using the clusterProfiler package [23,24], with adjusted p-value < 0.05. To assess direction-specific functional changes, DEPs from each pairwise comparison were further separated into upregulated and downregulated groups, and analyses were performed separately using the same enrichment workflow. Gene set enrichment analysis (GSEA) [25] was conducted using gseGO against GO:BP gene sets, with proteins ranked by limma t-statistics. Consensus clustering of DEPs was performed using the ConsensusClusterPlus package [26] (k=4, hierarchical clustering, Ward’s linkage, Euclidean distance, 500 resamplings). For each cluster, GO:BP and GO:CC enrichment, KEGG pathway analysis, and Reactome pathway analysis were performed using enrichGO, enrichKEGG (clusterProfiler), and enrichPathway (ReactomePA), respectively, with adjusted p-value < 0.05.

### Cell-Type-Specific Protein Signatures and Protein-Protein Interaction (PPI) Network Analysis

Direction-specific differential expression results from three pairwise comparisons were visualized using UpSet plots [27,28] and a circular heatmap based on upregulated and downregulated DEPs, enabling identification of shared, comparison-specific, and exact intersection subsets across comparisons, from which AS-EV, N-EV, MG-EV, and Glial-EV signature proteins were defined. Proteins significantly upregulated in both AS-EV vs N-EV and MG-EV vs N-EV but not significantly different between AS-EV and MG-EV were classified as a shared glial EV-enriched signature. Conversely, proteins significantly downregulated in both AS-EV vs N-EV and MG-EV vs N-EV but not in AS-EV vs MG-EV were defined as N-EV-enriched signatures. Proteins significantly upregulated in both AS-EV vs N-EV and AS-EV vs MG-EV with no significant difference between MG-EV and N-EV were defined as AS-EV-enriched signatures, and proteins significantly upregulated in both MG-EV vs N-EV and downregulated in AS-EV vs MG-EV but not in AS-EV vs N-EV were defined as MG-EV-enriched signatures. PPI networks of cell-type-specific DEPs were constructed using the Search Tool for the Retrieval of Interacting Genes (STRING) database [29] (interaction score ≥ 0.4). Network visualization was performed in R using the igraph and ggraph packages [30], with node size scaled by degree and edge width proportional to STRING combined score. Network layout was computed using the Fruchterman-Reingold algorithm. Molecular complex detection (MCODE) was performed using the MCODE algorithm in Cytoscape [31,32] to identify densely connected subclusters. Node-level topological parameters, including degree, betweenness centrality, closeness centrality, eigenvector centrality, and connected component membership, were calculated for each network and sub-network using the igraph package in R (degree, betweenness, closeness, eigen_centrality, and components functions, respectively). Betweenness and closeness centrality were normalized, with closeness computed within each connected component for networks containing more than one component.

## Results

### Characterization of Neuron-, Astrocyte-, and Microglia-Derived EVs from Cell Culture Medium

To characterize EVs derived from neuronal, astrocytic, and microglial cells, EVs were isolated from SH-SY5Y (N-EV), hAs (AS-EV), and HMC3 (MG-EV) cells by differential ultracentrifugation. EV morphology, size, particle concentration, and protein yield were assessed using TEM, NTA, and the BCA assay. TEM confirmed the characteristic cup-shaped bilayer membrane structure in all three EV populations (Figure 1A). NTA demonstrated that EVs from all three cell types showed a size distribution ranging from approximately 60 to 400 nm, with most particles concentrated between 80 and 150 nm (Figure 1B). No significant differences in average zeta potential were observed among the three EV groups (N-EV: −32.2 ± 4.1 mV; AS-EV: −33.9 ± 4.8 mV; MG-EV: −32.2 ± 4.3 mV, Figure 1C). Quantification revealed that N-EV particle concentration was significantly higher than that of both AS-EV (Figure 1D, 2.6 ± 0.1 vs 1.3 ± 0.1 particle concentration (×10¹¹ particles/mL) per 10⁷ cells, ****p < 0.0001) and MG-EV (2.6 ± 0.1 vs 0.75 ± 0.1 particle concentration (×10¹¹ particles/mL) per 10⁷ cells, ****p < 0.0001). In addition, AS-EV levels were significantly greater than those of MG-EV (*p < 0.05) (Figure 1D). Regarding EV protein amount, total EV protein yield per 10⁷ cells was significantly higher in N-EV (35.3 ± 8.4 µg) than in AS-EV (13.6 ± 1.0 µg) and MG-EV (11.4 ± 4.1 µg) (*p < 0.05; Figure 1E), consistent with N-EV having a higher particle concentration (Figure 1D). When normalized to particle number, however, protein content per 10⁹ EVs showed no significant differences across groups (N-EV: 1.3 ± 0.3 × 10⁵ pg, AS-EV: 1.1 ± 0.1 × 10⁵ pg, MG-EV: 1.6 ± 0.6 × 10⁵ pg; Figure 1F), indicating that protein loading per vesicle is maintained regardless of cell origin. Taken together, these results confirm successful EV isolation from cell media of SH-SY5Y, hAs, and HMC3 cells and demonstrate that AS-EV and MG-EV are secreted at lower quantities than N-EV, while their vesicle size and protein load per vesicle remain comparable across all three groups.

**Figure 1.**
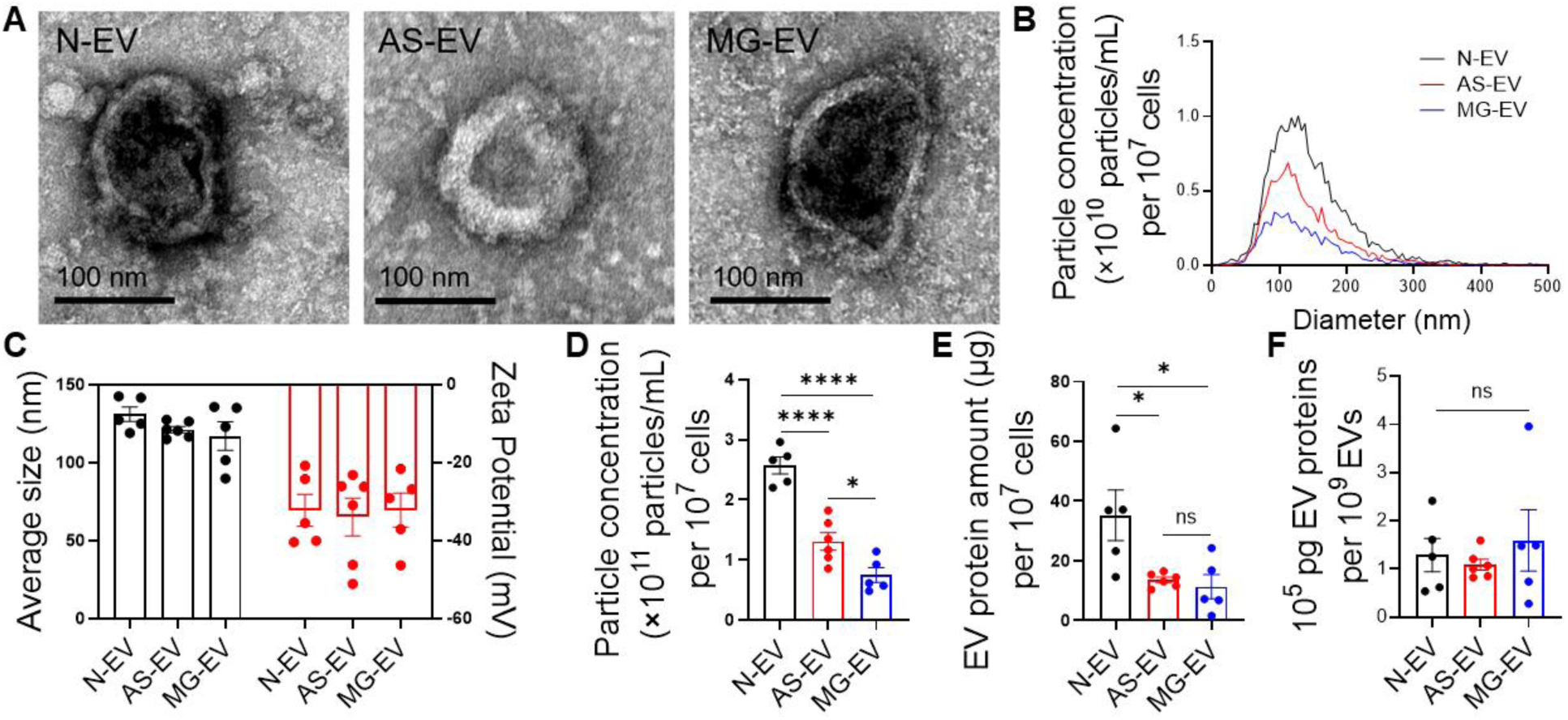
Characterization of neuron-, astrocyte-, and microglia-derived EVs. **A.** Representative TEM images of N-EV, AS-EV, and MG-EV. Scale bar = 100 nm. **B.** NTA of size distribution and particle concentration of N-EV, AS-EV, and MG-EV, normalized to 10⁷ cells. **C.** Comparison of average EV particle size and zeta potential across N-EV, AS-EV, and MG-EV groups. **D.** Quantification of EV particle concentration (×10¹¹ particles/mL) measured by NTA, normalized to 10⁷ cells. **E.** EV protein amount normalized to 10⁷ cells across N-EV, AS-EV, and MG-EV groups. **F.** EV protein content normalized to EV particle number across N-EV, AS-EV, and MG-EV groups. Data are presented as mean ± SEM. Statistical significance was determined by one-way ANOVA followed by Tukey’s post hoc multiple comparisons test; *p < 0.05, ****p < 0.0001. TEM, transmission electron microscopy; NTA, nanoparticle tracking analysis; EV, extracellular vesicle.

### Proteomic Profiles of N-EV, AS-EV, and MG-EV

Next, we performed 10-plex TMT-labeled LC-MS/MS to characterize the protein composition of N-EV, AS-EV, and MG-EV. A total of 3,022 proteins were identified across all three groups, of which 1,887 proteins passed quality filtering criteria and were included in downstream quantitative analysis. An omnibus F-test identified 922 proteins as DEPs across these three EV types (adjusted p-value < 0.01, |log₂FC| ≥ 1). PCA demonstrated distinct proteomic profiles across all three EV groups, with PC1 and PC2 accounting for 49% and 34.5% of the total variance, respectively (Figure 2A). Pairwise comparisons revealed 561 DEPs between AS-EV and N-EV, 608 DEPs between AS-EV and MG-EV, and 335 DEPs between MG-EV and N-EV. These differences were visualized by volcano plots, which showed 389 upregulated and 172 downregulated DEPs in AS-EV vs N-EV, 557 upregulated and 51 downregulated DEPs in AS-EV vs MG-EV, and 40 upregulated and 295 downregulated DEPs in MG-EV vs N-EV (Figure 2B). We next cross-referenced the 1,887 identified proteins against ExoCarta and Vesiclepedia, confirming that 1,662 (88.1%) and 1,783 (94.5%) of the identified proteins were present in each database, respectively (Figure 2C). Cross-referencing against published iPSC-derived neuron-, astrocyte-, and microglia-EV proteomes [15] further revealed that 73 (67%), 285 (75.4%), and 107 (54.3%) of proteins identified in each iPSC-derived EV dataset were detected in our N-EV, AS-EV, and MG-EV proteomes, respectively, while 36 (33%), 93 (24.6%), and 90 (45.7%) proteins reported in the iPSC-derived datasets were not detected in our corresponding cell-derived EVs (Figure 2D).

**Figure 2.**
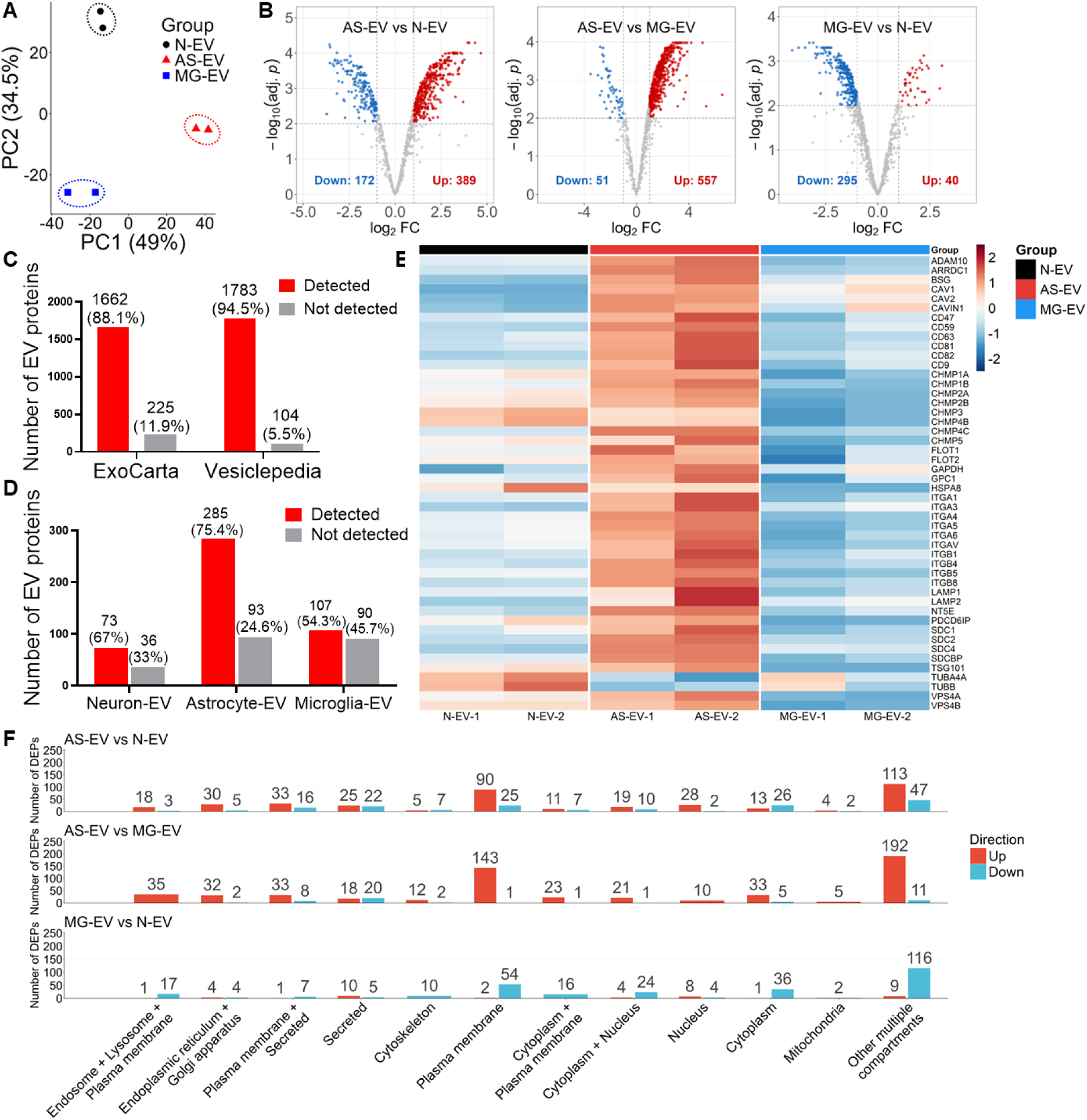
Comparative analysis of N-EV, AS-EV, and MG-EV proteomes. **A.** PCA of N-EV, AS-EV, and MG-EV proteomes (n = 2 per group) based on principal component 1 (49% variability) and principal component 2 (34.5% variability). **B.** Volcano plots showing protein distribution according to log₂FC and statistical significance (−log₁₀ adjusted p-value) in pairwise comparisons. Red and blue dots indicate upregulated and downregulated DEPs, respectively, whereas gray dots represent proteins with no significant differential expression. **C.** Overlap of identified EV proteins with human EV proteins curated in the ExoCarta and Vesiclepedia databases. **D.** Overlap of identified EV proteins with published iPSC-derived EV proteomes from iNeuron-EV, iAstrocyte-EV, and iMicroglia-EV [15]. **E.** Expression pattern of EV marker proteins described in the MISEV2023 guidelines [33] across N-EV, AS-EV, and MG-EV. Colors represent normalized protein abundance (z-score). **F.** Subcellular compartment distribution of DEPs in pairwise comparisons of N-EV, AS-EV, and MG-EV proteomes, annotated based on UniProt subcellular localization and GO terms, with unassigned proteins manually curated. Red and blue bars indicate upregulated and downregulated DEPs, respectively. PCA, principal component analysis; FC, fold change; DEPs, differentially expressed proteins; iPSC, induced pluripotent stem cell; MISEV, Minimal Information for Studies of Extracellular Vesicles; GO, Gene Ontology.

Canonical EV markers defined by Minimal Information for Studies of Extracellular Vesicles (MISEV) 2023 guidelines [33] were detected, including, but not limited to, transmembrane proteins such as tetraspanins (CD9, CD63, CD81, and CD82), integrins (ITGA/ITGB family), membrane-associated proteins (LAMP1 and LAMP2), and glycosylphosphatidylinositol (GPI)- or lipid-anchored proteins (GPC1, NT5E, and CD59), as well as cytosolic proteins including endosomal sorting complex required for transport (ESCRT) machinery components (TSG101, PDCD6IP, VPS4A/B, and CHMP family), flotillins (FLOT1/2), and caveolins (CAV1/2, and CAVIN1) (Figure 2E). In comparison, AS-EV showed relatively higher abundance of these EV marker proteins than N-EV and MG-EV, while MG-EV displayed relatively lower expression of most markers. Subcellular compartment analysis of DEPs from pairwise comparisons revealed distinct protein distribution patterns among AS-EV, MG-EV, and N-EV proteomes (Figure 2F). Compared with N-EV and MG-EV, AS-EV showed enrichment of plasma membrane-, secreted-, and membrane-associated secreted proteins, suggesting differences in extracellular communication-related cargo loading (Figure 2F). Notably, endosome- and lysosome-associated proteins were also upregulated in AS-EV compared to N-EV and MG-EV, potentially reflecting active vesicular trafficking in astrocytes. Furthermore, enrichment of endoplasmic reticulum- and Golgi-associated proteins in AS-EV suggested enhanced secretory and protein-processing activity in astrocytes. In contrast, comparison of MG-EV vs N-EV showed a markedly different pattern, with a predominance of downregulated proteins across most compartments. The greatest protein abundance differences were observed between AS-EV and MG-EV, particularly in plasma membrane-associated proteins and proteins assigned to multiple cellular compartments (Figure 2F). Lastly, across all three comparisons, “other multiple compartments” proteins accounted for the largest proportion, indicating the complexity of EV protein subcellular origin (Figure 2F). Taken together, these findings demonstrate cell-type-specific heterogeneity in EV protein composition and subcellular cargo distribution.

### Functional Characterization of AS-EV, MG-EV, and N-EV by Pathway Enrichment Analysis and Consensus Clustering

GSEA based on GO:BP gene sets (Figure 3A), together with GO:BP, GO:CC, and KEGG pathway enrichment analyses (Figure 3B), was performed across the three pairwise comparisons to characterize functional differences among the three EV types. GSEA enrichment plots for selected GO:BP gene sets are presented in Supplemental Figure S1A-F. GO:BP, GO:CC, and KEGG enrichment analyses were also performed separately on upregulated (Supplemental Figure S2) and downregulated DEPs (Supplemental Figure S3). Cell adhesion, extracellular matrix (ECM) interaction, and host interaction and symbiont interaction represented the most consistent signature of AS-EV, with enriched focal adhesion, cell-substrate junction, ECM-receptor interaction, integrin signaling, cell adhesion molecule (CAM) interaction, and host and symbiont interaction (GSEA, GO:BP, GO:CC, KEGG, up GO:BP, up GO:CC and up KEGG) relative to both N-EV and MG-EV. Endosomal trafficking and multivesicular body (MVB) biogenesis were a common feature of AS-EV and N-EV compared to MG-EV, with MVB organization and assembly, ubiquitin (Ub)-dependent protein catabolic process via the MVB sorting pathway, endosome transport via multivesicular body sorting pathway, late endosome to vacuole transport, and viral budding via host ESCRT complex enriched in both EV types (GSEA, GO:BP, up GO:BP and down GO:BP). In contrast, MG-EV was enriched in chromatin organization and epigenetic regulatory processes relative to N-EV, with significantly upregulated pathways including nucleosome assembly, heterochromatin formation, protein-DNA complex organization, and epigenetic regulation of gene expression identified by GSEA. Relative to AS-EV, MG-EV was enriched in immune, cytotoxic, and coagulation-related processes, including killing or disruption of cells of another organism and blood coagulation, as well as protein metabolism-related processes, including tRNA aminoacylation, amino acid activation, and protein processing (GSEA and down GO:BP). In contrast to AS-EV, N-EV was enriched in complement and coagulation processes, as well as neuronal morphology-related pathways including axon and neuronal projection guidance, as indicated by down GO:BP and down KEGG. Additionally, N-EV was distinguished from MG-EV by enrichment of endocytic and membrane-associated components, including cell leading edge, ruffle, cell projection membrane, and ESCRT complex (GO:CC and down GO:CC), as well as diverse signaling pathways including endocytosis, immunoglobulin superfamily (IgSF) CAM signaling, Ras, Rap1, and chemokine signaling (KEGG and down KEGG), and actin filament polymerization (GSEA), suggesting that N-EV cargo is associated with membrane dynamics and receptor-mediated signal transduction.

**Figure 3.**
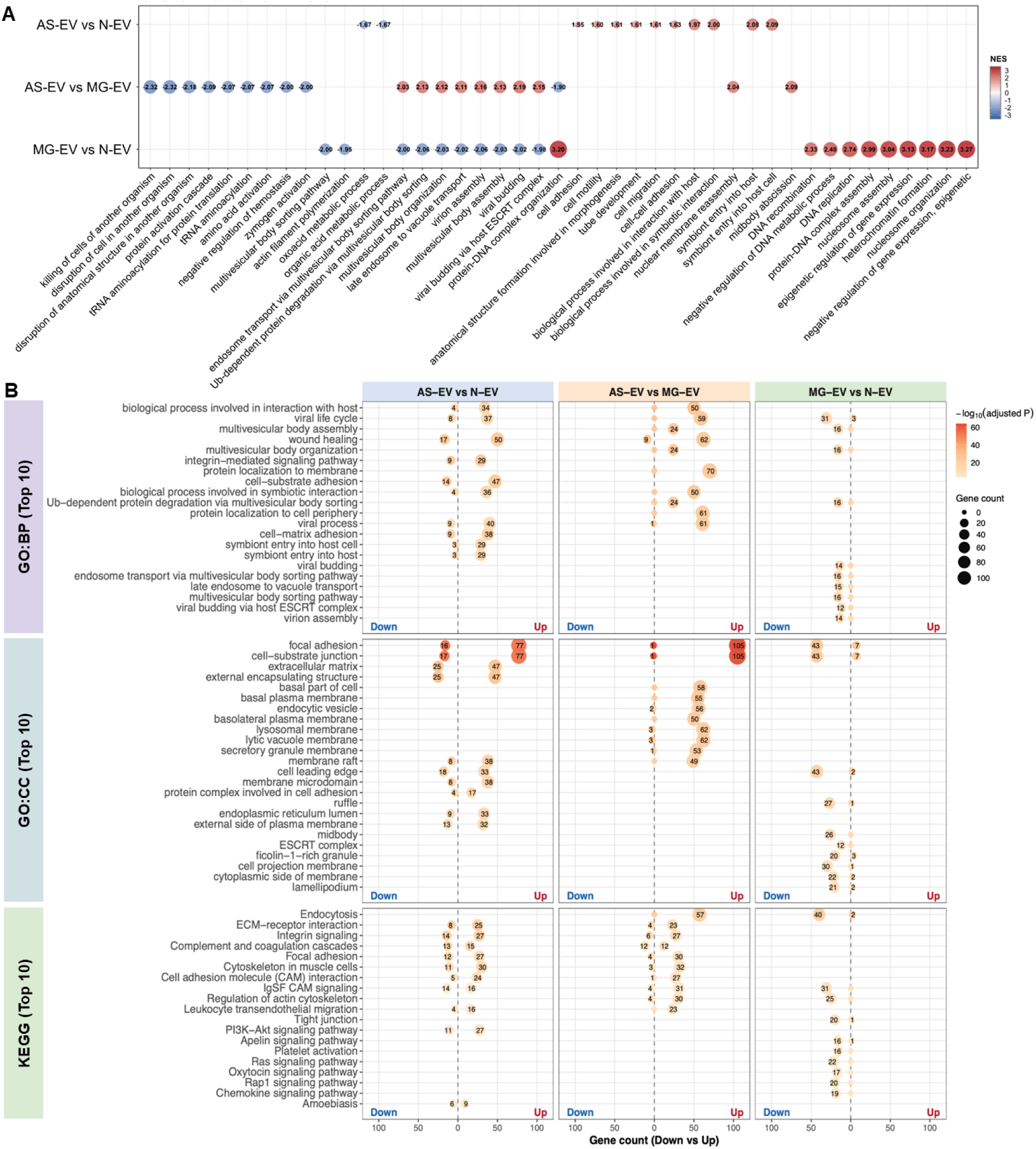
Functional enrichment analysis across EV groups. **A.** GSEA bubble plot showing NES of significantly enriched GO:BP terms (adjusted p-value < 0.05) across the three pairwise comparisons. Up to the top 10 positively enriched (red, NES > 0) and 10 negatively enriched (blue, NES < 0) terms are shown for each comparison. NES values are shown within each bubble. **B.** Dot plots showing up to the top 10 enriched GO:BP terms, GO:CC terms, and KEGG pathways (adjusted p-value < 0.05) for upregulated and downregulated DEPs in the AS-EV vs N-EV, AS-EV vs MG-EV, and MG-EV vs N-EV comparisons. Dot size represents gene count, and color indicates statistical significance (−log₁₀ adjusted p-value). GSEA, gene set enrichment analysis; NES, normalized enrichment score; GO:BP, GO biological process; GO:CC, GO cellular component; KEGG, Kyoto Encyclopedia of Genes and Genomes.

We further performed consensus clustering (k=4) to identify the heterogeneity of DEP expression patterns and their corresponding functions (Figure 4 and Supplemental Figure S4). The row-scaled z-score heatmap confirmed the different expression profiles across these four clusters, with cluster 1 co-enriched in N-EV and MG-EV, cluster 2 co-enriched in N-EV and AS-EV, cluster 3 specifically enriched in AS-EV, and cluster 4 specifically enriched in N-EV (Supplemental Figure S4A). The distribution of upregulated and downregulated DEPs across pairwise comparisons is shown in Supplemental Figure S4B. GO:CC, KEGG, and Reactome [34] enrichment analyses of the four clusters are presented in Supplemental Figure S4C, S4D, and S4E, respectively. Cluster 1 (N-EV ≈ MG-EV > AS-EV, n=96) was enriched in complement activation, coagulation, and humoral immune responses, consistent with the complement and coagulation cascades identified across the pairwise comparisons against AS-EV (Figure 4A and Supplemental Figure S3, S4D-E). Cluster 2 (N-EV ≈ AS-EV > MG-EV, n=213) recapitulated the MVB biogenesis signature (Figure 4A and Supplemental Figure S4C-E) identified by GSEA and GO:BP (Figure 3A–B, Supplemental Figure S2, Supplemental Figure S3) in the pairwise comparisons of AS-EV and N-EV against MG-EV, with enrichment in endosome organization, MVB assembly, and viral budding via the host ESCRT complex. The AS-EV-specific cluster 3 (n=403) was dominated by cell adhesion, cell surface receptor signaling, and host and symbiont interaction terms (Figure 4A and Supplemental Figure S4C-E), a pattern also captured by GSEA, GO:CC, and KEGG (Figure 3A-B and Supplemental Figure S2) in the pairwise comparisons of AS-EV against both N-EV and MG-EV. Cluster 4 (N-EV-enriched, n=210) was enriched in actin filament polymerization, regulation of actin filament polymerization, regulation of supramolecular fiber organization, and protein polymerization (Figure 4A). The actin filament polymerization result was consistent with GSEA in N-EV relative to MG-EV (Figure 3A) and was further supported by related axon guidance enrichment among downregulated DEPs in AS-EV vs N-EV (Supplemental Figure S3). Cluster 4 was also enriched in other biological processes, including cellular response to hormone stimulus, response to hormone, and G protein-coupled receptor signaling pathway (Figure 4A and Supplemental Figure S4D-E).

**Figure 4.**
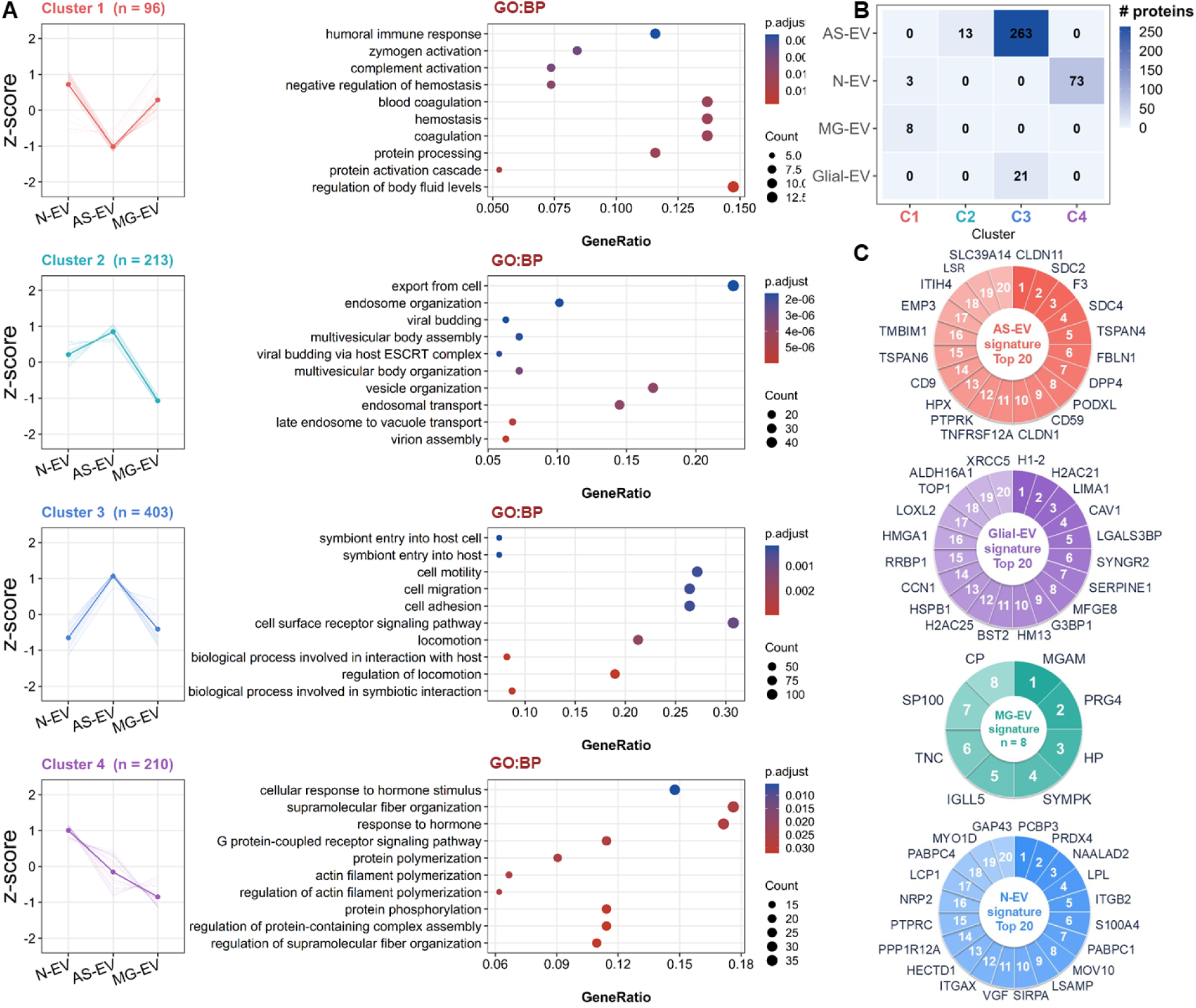
Consensus clustering and EV cell-type signature protein analysis. **A.** Consensus clustering (k=4) of common DEPs identified by F-test using ConsensusClusterPlus with hierarchical clustering. Left panels show row-scaled z-score expression profiles for each cluster (thin lines represent individual proteins; thick lines represent the cluster mean). Right panels show up to the top 10 enriched GO:BP terms per cluster (adjusted p-value < 0.05). Bubble size represents gene count, and color indicates adjusted p-value. **B.** Heatmap showing the distribution of independently defined EV cell-type signature proteins across the four consensus clusters identified in panel A (C1–C4). Color intensity represents protein count. **C.** Circular ranked plots of EV cell-type signature proteins for AS-EV, Glial-EV, MG-EV, and N-EV. Numbered sectors indicate the relative ranking of proteins within each signature, integrating differential effect size and statistical significance across the relevant pairwise comparisons. The top 20 signature proteins are shown for AS-EV, Glial-EV, and N-EV, whereas all 8 MG-EV signature proteins are displayed.

### Identification of EV Cell-Type Signature Proteins and Functional Interaction Networks

Cell-type-specific EV cargo was further investigated by defining EV signature proteins and mapping them onto DEP-based consensus clustering. Four groups of EV signature proteins were defined from shared DEPs with the F-test (adjusted p-value < 0.01, |log₂FC| ≥ 1). Direction-specific UpSet plot analysis revealed shared and comparison-specific DEPs across EV groups (Supplemental Figure S5). The UpSet plot summarized exact intersection patterns derived from the three pairwise comparisons, with DEPs separated into up- and downregulated sets for each comparison, distinguishing proteins that are comparison-specific from those shared across multiple comparisons. Two-dot intersections can show shared signatures, such as proteins increased in both AS-EV and MG-EV relative to N-EV. These intersection-defined signatures were then visualized in the circular heatmap to show their relative abundance patterns across N-EV, AS-EV, and MG-EV. Exact intersection patterns identified 276 AS-EV signature proteins, 76 N-EV signature proteins, 8 MG-EV signature proteins, and 21 Glial-EV signature proteins, all of which displayed distinct abundance patterns as visualized in the circular heatmap (Supplemental Figure S5). EV signature proteins were then mapped onto the four consensus clusters (Figure 4B). AS-EV signature proteins were predominantly assigned to cluster 3 (n=263), with smaller subsets in cluster 2 (n=13), consistent with the AS-EV-enriched expression patterns of these clusters and their functions in cell adhesion, interaction, and endosomal trafficking. N-EV signature proteins were predominantly assigned to cluster 4 (n=73), with a small subset in cluster 1 (n=3), recapitulating the function of cluster 4 in actin cytoskeleton organization and biological processes. MG-EV signature proteins were exclusively assigned to cluster 1 (n=8), while Glial-EV signature proteins were exclusively assigned to cluster 3 (n=21), reflecting the shared AS-EV and MG-EV component captured by this cluster. Expression of EV cell-type signature proteins was visualized in a circular plot displaying up to the top 20 proteins per signature (Figure 4C). PPI network was performed to identify hub proteins and functional interaction modules within each EV cell type signature proteins mapping on corresponding consensus clustering (Figure 5A-D).

**Figure 5.**
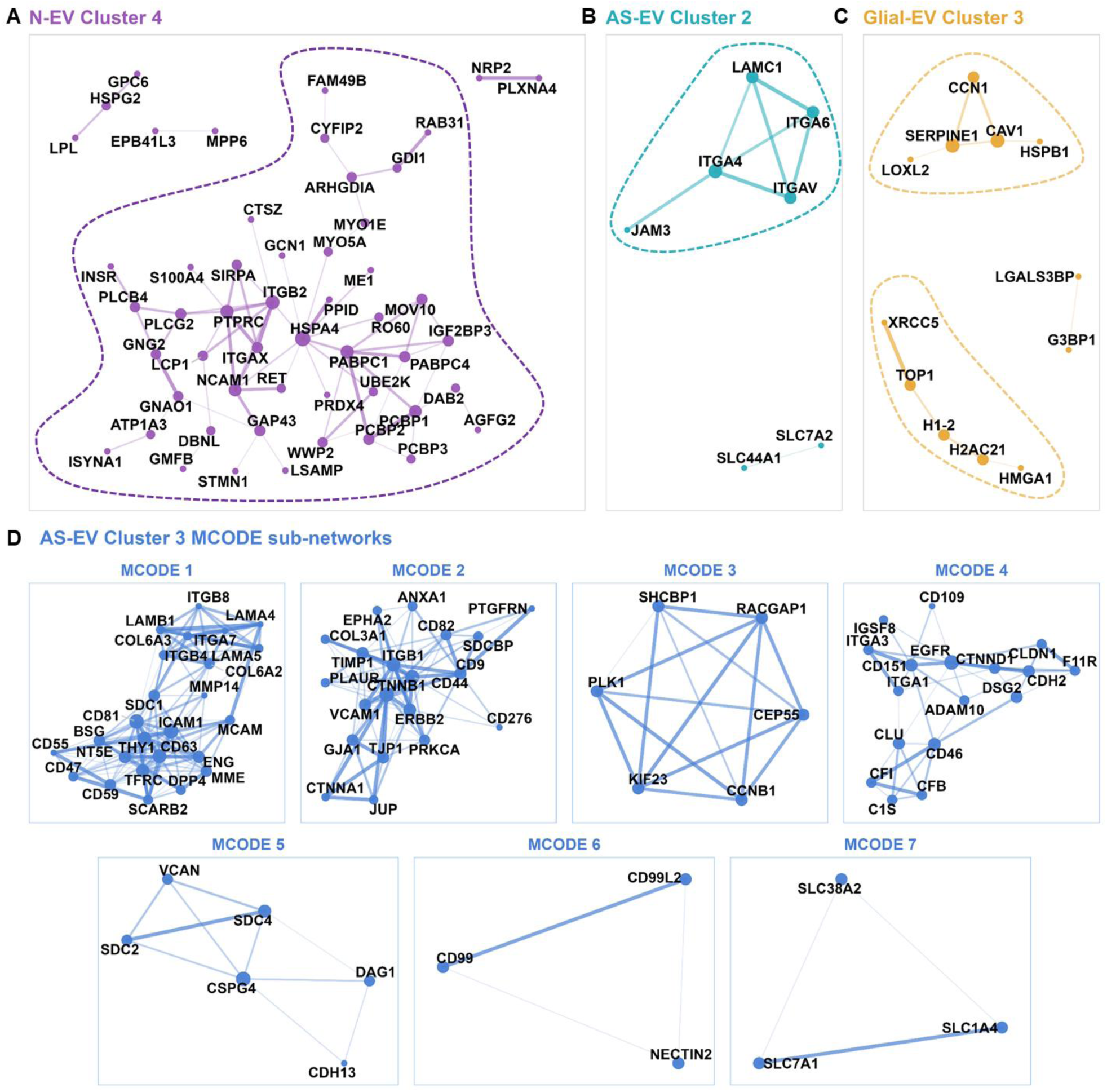
PPI networks of EV cell-type signature proteins across consensus clusters. For each cluster, signature proteins assigned to that cluster were submitted to the STRING database (combined score ≥ 0.4) for PPI network analysis. Node size reflects degree, and edge width reflects interaction confidence. **A.** N-EV signature proteins in Cluster 4. **B.** AS-EV signature proteins in Cluster 2. **C.** Glial-EV signature proteins in Cluster 3. **D.** AS-EV signature proteins in Cluster 3; seven MCODE sub-networks (MCODE 1–7) are shown individually. PPI, protein-protein interaction; STRING, Search Tool for the Retrieval of Interacting Genes; MCODE, Molecular Complex Detection.

Of the 73 N-EV signature proteins assigned to cluster 4, 52 were mapped in STRING and included in the PPI analysis. The N-EV cluster 4 network consisted of one major connected component containing 45 of the 52 proteins (Figure 5A). The network comprised proteins associated with several biological functions, including immune-associated proteins (e.g., ITGB2, PTPRC, and ITGAX), RNA-binding proteins (e.g., PABPC1, PCBP1, and PCBP2), cytoskeletal and vesicle-trafficking proteins (e.g., ARHGDIA, GDI1, and RAB31), and neuronal proteins (e.g., NCAM1, GAP43, and STMN1). HSPA4 represented the highest-degree node linking multiple functional modules within the network. The major connected component of the AS-EV cluster 2 network represented an integrin-associated cell adhesion and ECM network, with ITGAV, ITGA4, and ITGA6 interconnected with the ECM protein LAMC1 and the adhesion molecule JAM3 (Figure 5B). Although cluster 2 was globally enriched for endosomal trafficking and MVB-associated proteins, this PPI subnetwork highlighted an additional adhesion-related module assigned to this cluster. The Glial-EV PPI network was organized into three connected components (Figure 5C). The two major networks were enriched for extracellular matrix-associated proteins (e.g., CAV1, CCN1, SERPINE1, and LOXL2) and chromatin-associated proteins (e.g., TOP1, XRCC5, H1-2, H2AC21, and HMGA1). The AS-EV cluster 3 network was the largest and most complex PPI network and was further subclustered into seven functional modules using the MCODE algorithm in Cytoscape (Figure 5D). MCODE analysis identified a tetraspanin (CD63/CD81)-associated surface protein module with integrin-laminin adhesion components (MCODE 1), a tetraspanin-associated EV protein and cell-cell adhesion module (MCODE 2), a mitotic cell cycle module (MCODE 3), a cell adhesion, receptor signaling, and complement regulation module (MCODE 4), a proteoglycan-rich ECM module (MCODE 5), a cell-cell adhesion module (MCODE 6), and an amino acid transport module (MCODE 7).

## Discussion

In this study, we performed TMT-based quantitative proteomic profiling of EVs isolated from representative human neuronal-like (SH-SY5Y), astrocytic (hAs), and microglial (HMC3) cells under baseline conditions to enable a direct side-by-side comparison of their protein cargo. Comparative proteomics revealed cell-type-specific heterogeneity in EV protein cargo and suggested distinct functional roles of EVs in intercellular communication. These proteomic datasets serve as a baseline reference for future disease-state comparisons and EV-based drug delivery and therapeutic applications.

Cell-type heterogeneity was first evident in EV secretion. N-EV showed the highest particle concentrations, followed by AS-EV and MG-EV, and total EV protein yield was also highest in N-EV, with no significant difference between AS-EV and MG-EV. Despite these differences in EV output, average EV size, zeta potential, and protein content per vesicle were comparable across the three cell types, suggesting that cellular origin primarily influenced EV production yield rather than the measured biophysical properties or average protein loading of individual vesicles. Previous studies have reported different patterns of EV secretion across CNS cell types. For example, astrocyte-derived EVs exhibited higher particle concentrations than neuron-derived EVs in conditioned media from human astrocytes (Gibco/Thermo Fisher Scientific) and human neurons (ScienCell Research Laboratories) [35] as well as in EVs isolated by immunocapture from C57BL/6 mouse brain tissue, where microglia-derived EVs showed the lowest yield [14]. The low EV output observed in microglia in the latter study is consistent with their role as resident immune cells, whose EV release can be modulated by immune activation and inflammatory stimuli [36]. In contrast, astrocytes may constitutively secrete EVs as part of their homeostatic support functions, including metabolic and trophic maintenance of neurons [37], while the highest EV secretion observed in N-EV may reflect the active intercellular communication demands of neurons, which may rely on vesicle-mediated signaling to coordinate synaptic function and neural circuit activity [38]. Nevertheless, these functional roles do not necessarily predict the relative magnitude of EV secretion under specific experimental conditions. In the present study, N-EV production exceeded AS-EV production, differing from the trends reported in previous studies. This discrepancy suggests that relative EV output is not necessarily conserved across cellular models and may depend on cell characteristics, culture conditions, and experimental approaches to EV isolation and quantification. Thus, the observed differences in EV production likely reflect model- and context-dependent secretory properties rather than a universal hierarchy of EV secretion among CNS cell types.

Proteomic profiling further revealed cell-type-specific heterogeneity in EV protein composition, with PCA demonstrating clear separation among N-EV, AS-EV, and MG-EV. Most identified proteins were represented in established EV databases, including ExoCarta and Vesiclepedia, supporting the vesicular identity of the isolated fractions. Canonical EV proteins, including tetraspanins, ESCRT-associated proteins, and integrins, were detected in all three EV groups but showed the highest abundance in AS-EV and the lowest in MG-EV. This pattern was consistently reflected across differential proteomic analyses, where volcano plots, direction-specific circular heatmaps, and subcellular compartment classification demonstrated a graded pattern of significantly upregulated DEPs (AS-EV > N-EV > MG-EV), indicating that these cell-type differences extend beyond canonical EV markers to the broader proteomic composition. In particular, this graded pattern was consistently observed across most subcellular compartments rather than being restricted to a particular protein category. Such changes suggest that the observed differences reflect broad cell-type-dependent variation in EV cargo composition, rather than selective enrichment or depletion of proteins originating from individual cellular compartments. The predominance of AS-EV-enriched DEPs together with the relatively small number of MG-EV-enriched DEPs suggests that the MG-EV proteome is comparatively less distinct under basal conditions. Interestingly, these compositional differences were only partially mirrored by EV production. MG-EV showed the lowest values, consistent with their comparatively less distinct proteomic profile. In contrast, although AS-EV was produced at lower levels than N-EV, they exhibited the highest abundance of canonical EV proteins and the largest number of enriched DEPs in pairwise comparisons. These findings suggest that the quantity of EVs released and their protein cargo represent distinct but complementary features of CNS EV biology [39,40]. One possible explanation is that astrocytes, neurons, and microglia differ intrinsically in EV cargo selection under basal conditions. Astrocytes may selectively enrich a broader repertoire of proteins within their EVs despite producing fewer vesicles than neurons, potentially reflecting a greater contribution of EV populations enriched in canonical endolysosomal and membrane-associated EV machinery. This interpretation is consistent with previous neural cell EV proteomic studies showing cell-type-specific differences in EV protein composition [15] and with recent evidence that astrocyte-derived EVs exhibit broader selective protein enrichment relative to their parent cells [35]. In contrast, HMC3-derived EVs showed lower representation of canonical EV-associated proteins, which may reflect a less distinct basal EV proteome in this model.

The integration of GSEA, GO, and KEGG pathway enrichment analyses, together with consensus clustering and PPI network analysis, demonstrated distinct functional differences among EVs derived from neurons, astrocytes, and microglia. AS-EV exhibited a protein profile consistent with the homeostatic functions of astrocytes in the CNS, including regulation of the extracellular environment, metabolic and structural support, and communication with neurons and other glial cells [41,42]. Consistent with these roles, AS-EV-enriched proteins converged on ECM organization, cell adhesion, integrin signaling, and vesicular trafficking, indicating that AS-EV cargo is molecularly equipped to support extracellular remodeling and cell-cell interactions within the neural microenvironment, extending the supportive functions of astrocytes beyond the parent cell. Several protein classes enriched in the AS-EV proteome have previously been reported to localize on the EV surface and participate in EV-recipient cell interactions, providing a mechanistic basis for this extended function. Among these, integrins are well-established regulators of EV docking, target-cell recognition, and downstream signaling, whereas cell adhesion molecules (including members of the CAM family) contribute to stable EV-cell interactions and recipient-cell targeting. Proteoglycans and extracellular matrix proteins, including laminins, have also been implicated in facilitating EV docking by providing adhesive interfaces and cooperating with integrins during the initial interaction with recipient cells [43,44]. Importantly, PPI analysis of AS-EV signature proteins further reinforced this finding by demonstrating that these cell-type-specific proteins formed highly interconnected interaction networks centered on ECM interaction, integrin signaling, proteoglycan-rich ECM, and cell junction organization. Consistent with our findings, a proteomic analysis of human astrocyte-derived EVs by Chun et al. (2021) found that, relative to a generic EV protein background, astrocyte-derived EV proteins were significantly enriched for integrin binding, ECM structural constituent, and laminin binding among molecular functions, with ECM organization emerging as the top enriched biological process [19]. This integrin enrichment has been independently corroborated at the single-vesicle level. Immuno-TEM has shown that 71.6% of human adult astrocyte-derived EVs are positive for integrin β1 [45]. Quantitative comparison further showed that astrocyte-derived EVs express markedly higher levels of integrin α than neuron-derived EVs [20]. Overall, the coordinated enrichment and organization of these adhesion- and matrix-associated proteins suggest that AS-EV package functionally related cargo specialized for mediating cell-cell and cell-matrix interactions, thereby promoting productive interactions with recipient cells, facilitating EV uptake, and supporting structural integrity and astrocyte-mediated intercellular communication within the CNS. These molecular characteristics may also contribute to the growing interest in astrocyte-derived EVs as CNS drug delivery vehicles, as intranasally administered astrocyte-derived EVs loaded with ultrasmall superparamagnetic iron oxide nanoparticles have been shown to distribute into the brain, supporting their feasibility as a CNS-targeted therapy [46].

In contrast, N-EV displayed neuron-specific processes related to connectivity and structural remodeling rather than extracellular matrix regulation. N-EV-enriched proteins were associated with axon and neuronal projection guidance, actin cytoskeleton organization, and receptor-mediated signaling pathways including endocytosis, Ras, Rap1, and chemokine signaling, consistent with roles for neurons in neuronal and synaptic plasticity and cytoskeleton remodeling [47–53]. Proteins involved in cytoskeletal dynamics and vesicular trafficking (e.g., ARHGDIA, GDI1, and RAB31), together with neuronal development- and axon-remodeling-associated proteins (e.g., NCAM1, GAP43, and STMN1), further support a role for N-EV cargo in facilitating axonal and dendritic remodeling and neuron-to-neuron signaling relevant to circuit maintenance. These findings align with prior reports that neuron-derived EVs carry synaptic proteins and actively modulate synaptic transmission and plasticity [54]. In addition, PPI analysis of N-EV signature proteins also identified a subnetwork containing RNA-binding proteins (e.g., PABPC1, PCBP1, and PCBP2), suggesting a potential contribution of N-EV cargo to post-transcriptional regulation. Although the present proteomic analysis was performed on bulk EVs and therefore cannot determine whether these proteins coexist within individual vesicles, the integration of neuronal, RNA-binding, vesicle-trafficking, and immune-associated proteins into a single PPI network suggests that N-EV collectively carries coordinated molecular programs supporting neuronal homeostasis and intercellular communication. Furthermore, the shared enrichment of endosomal and multivesicular body-associated proteins in AS-EV and N-EV, relative to MG-EV, may reflect a greater contribution of the endosomal pathway to EV biogenesis and selective cargo sorting before secretion.

Relative to AS-EV, N-EV and MG-EV shared enrichment of complement and coagulation cascade-related proteins. Complement proteins are increasingly recognized to have functions within the CNS beyond their canonical immune roles, and neuronal cells have been shown to express complement components [55]. Human neuronal cells, including the SH-SY5Y cell line, can constitutively synthesize complement components such as C1q, C3, and C4, indicating that neurons can serve as a local source of complement proteins within the CNS under basal conditions [56]. Our findings showed that N-EV were enriched in complement-related proteins, raising the possibility that these vesicles contribute to localized complement-related signaling within the neural microenvironment. MG-EV similarly showed enrichment of complement-associated proteins, consistent with recent evidence that microglia-derived EVs carry complement proteins involved in synaptic remodeling [57]. PPI analysis of the N-EV network additionally identified a network enriched for leukocyte-associated proteins including PTPRC, ITGB2, and ITGAX, whose interpretation requires further validation [58,59]. In addition, MG-EV displayed distinct cytotoxic and immune-effector pathway enrichment relative to AS-EV that was not observed in N-EV, consistent with microglia’s canonical immune surveillance role [60]. These findings suggest that although N-EV and MG-EV share complement-associated features, they exhibit distinct immune-related cargo profiles that are likely to reflect their cell-type-specific roles in maintaining CNS homeostasis under physiological conditions.

Relative to N-EV, however, MG-EV was additionally enriched for nucleosome assembly, heterochromatin formation, protein-DNA complex organization, and other chromatin-associated pathways, suggesting that microglia package chromatin-associated proteins into EVs. Chromatin-associated proteins, particularly histones, have been consistently identified in EV proteomic studies [61]. Although the functional significance of this cargo remains unclear, a recent study has demonstrated that EVs can carry nuclear proteins, including histones, and that selective sorting of nuclear proteins into EVs is a regulated process rather than a consequence of passive cellular leakage [62]. Because only eight MG-EV signature proteins were identified, PPI network analysis was not performed, as the limited number of proteins precluded meaningful network construction. Interestingly, this chromatin-associated feature was not exclusive to MG-EV. We identified a Glial-EV signature shared by AS-EV and MG-EV but absent from N-EV, indicating that packaging of chromatin-associated and epigenetic regulatory proteins may be a shared feature of glial-derived EVs in these models rather than a microglia-specific characteristic. PPI analysis of the Glial-EV network further resolved two major functionally distinct modules: an extracellular matrix remodeling module (CAV1, SERPINE1, CCN1, and LOXL2) and a chromatin regulation module (H1-2, H2AC21, HMGA1, TOP1, and XRCC5), suggesting that Glial-EV cargo may support two functional programs, each consistent with roles described for AS-EV and MG-EV.

Taken together, the distinct proteomic signatures of N-EV, AS-EV, and MG-EV recapitulate the specialized physiological roles of their parental cell types: neuronal connectivity and structural remodeling, astrocytic extracellular support and cellular interaction, and microglial immune surveillance. In addition, this study identified molecular features extending beyond these canonical functions, including complement-associated cargo in both N-EV and MG-EV and an unexpected chromatin-regulatory signature shared between AS-EV and MG-EV, highlighting potential roles for CNS-derived EVs in immune signaling and chromatin-associated intercellular communication. Beyond reflecting parental cell identity, these cargo signatures suggest that EVs have distinct molecular characteristics that may contribute to cell-type-specific functions within the CNS.

We recognize several limitations of our study. First, each EV type was profiled with two biological replicates, which may limit statistical power and the precision of variance estimates [63]. We sought to partly offset this by analyzing all samples in a single TMT 10-plex experiment [64], using empirical Bayes-moderated statistics [65,66] with stringent thresholds that are, if anything, conservative given TMT ratio compression [67], and emphasizing pathway-level patterns over individual proteins. Even so, we view these data as a preliminary baseline, and smaller differences should be interpreted cautiously and confirmed in independent preparations. Second, EVs were derived from cultured human cell models, namely SH-SY5Y neuroblastoma cells, SV40-immortalized HMC3 microglia, and primary human astrocytes, rather than CNS cells in vivo or brain tissue, and cell lines may not fully retain the functions of their primary counterparts [68]. These models allowed all three EV types to be generated and processed in parallel under identical conditions, avoiding the donor, regional, and cell-mixture variability of tissue-derived EVs. The overlap with iPSC-derived neural EV proteomes [15] (Figure 2D) is encouraging but does not replace validation in primary and in vivo-derived EVs. Third, differential ultracentrifugation does not resolve EV subtypes and may co-isolate non-vesicular material. We chose this widely used method to facilitate comparison with existing datasets, applied it identically across cell types, and use the operational term “EVs” rather than “exosomes”. Whether signature proteins are shared across subtypes will require subtype-resolved approaches [69,70]. Finally, EV cargo functions were inferred from enrichment analyses rather than experimentally validated. To reduce reliance on any single method, we combined gene set enrichment and over-representation analyses [71] with consensus clustering and interaction networks, and focused on themes consistent across analyses and with prior literature. These inferences are best regarded as hypotheses.

Each of these limitations points to a concrete next step. Extending this framework to primary and iPSC-derived neural cells, to activated, aged, and disease-relevant states, and to EVs from brain tissue and biofluids will test how well these baseline signatures generalize in vivo. Subtype-resolved isolation and single-vesicle approaches could clarify whether signature proteins co-occur within individual vesicles, and uptake and functional assays in recipient cells will be needed to define the roles of candidate cargo, including complement- and chromatin-associated proteins. Integrating lipidomic and RNA profiles would further extend this molecular description. As these layers accumulate, cell-type-specific signatures may help attribute the cellular origin of EVs in cerebrospinal fluid and blood and guide the rational selection, and ultimately the engineering, of EVs for targeted delivery to the brain. By defining what each neural cell type releases under baseline conditions, this study provides a reference against which disease-associated changes in EV cargo can be interpreted and EV-based applications developed.

## Author Contributions

C. H. L. and J. Z. conceptualization; X. C. methodology; X. C. investigation; X. C. formal analysis; X. C. data curation; X. C. visualization; X. C., C. H. L., and J. Z. writing – original draft; C. H. L. and J. Z. writing – review and editing; C. H. L. and J. Z. supervision; C. H. L. and J. Z. project administration; C. H. L. and J. Z. funding acquisition; C. H. L. and J. Z. resources.

## Acknowledgements

The authors thank Dr. Ebbing de Jong of the Proteomics & Mass Spectrometry Core Facility at SUNY Upstate Medical University for performing the LC-MS/MS proteomics analysis and for providing the sample preparation and data acquisition protocols, and Benjamin Zink, TEM Core Technical Manager from the Upstate Medical University TEM Core, for TEM sample preparation and imaging.

## Data Availability Statement

The data that support the findings of this study are available from the corresponding author upon reasonable request.

