## Supplementary Figures for "Comparative Proteomics Reveals Cell-Type-Specific Heterogeneity of Human Neural Cell-Derived Extracellular Vesicles"

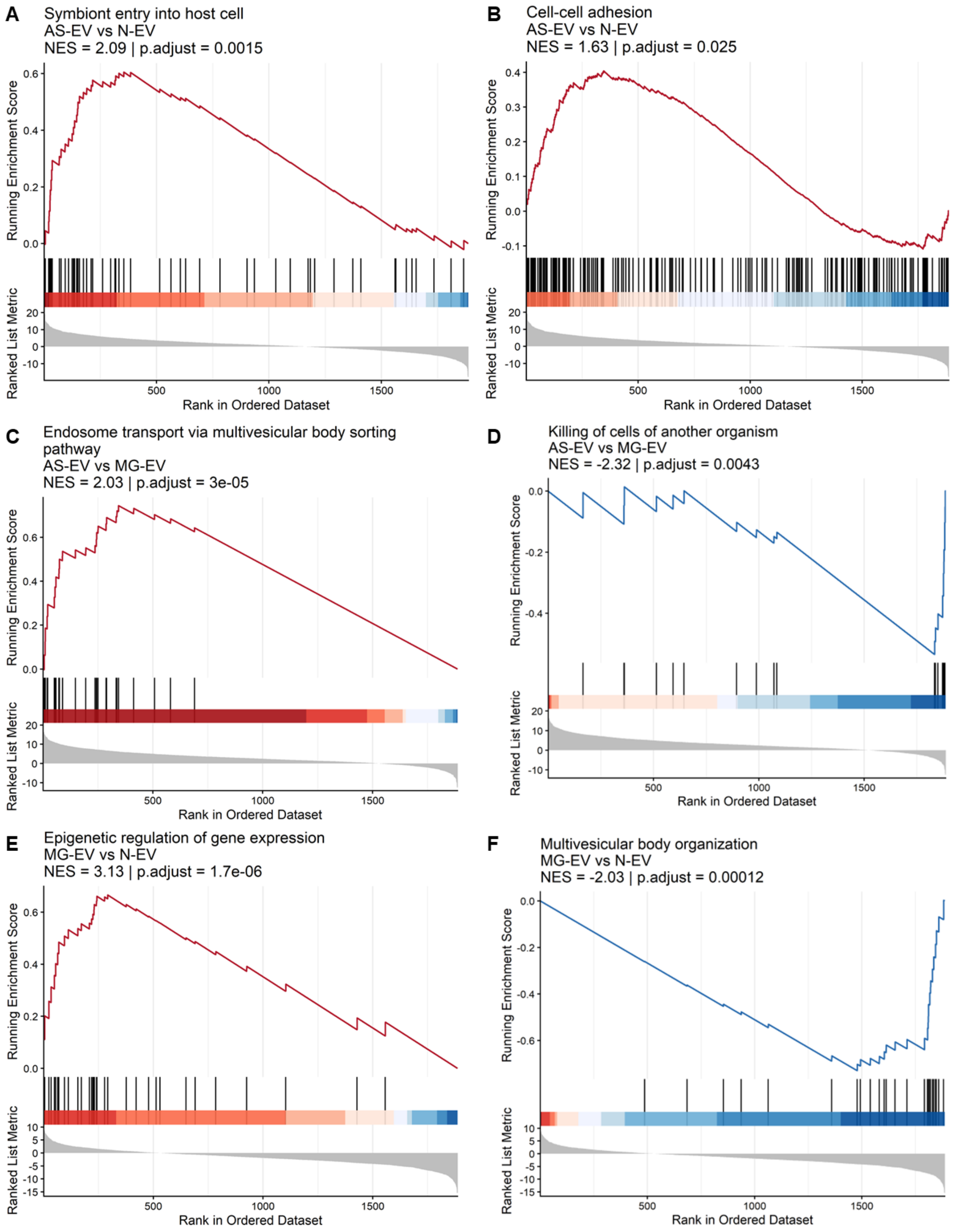


**Supplemental Figure S1** Representative GSEA enrichment plots of selected GO gene sets across EV pairwise comparisons. GSEA was performed for the three pairwise comparisons among N-EV, AS-EV, and MG-EV. **A.** Symbiont entry into host cell, AS-EV vs N-EV. **B.** Cell-cell adhesion, AS-

EV vs N-EV. **C.** Endosome transport via multivesicular body sorting pathway, AS-EV vs MG-EV. **D.** Killing of cells of another organism, AS-EV vs MG-EV. **E.** Epigenetic regulation of gene expression, MG-EV vs N-EV. **F.** Multivesicular body organization, MG-EV vs N-EV. NES and adjusted p-values are indicated in each panel.

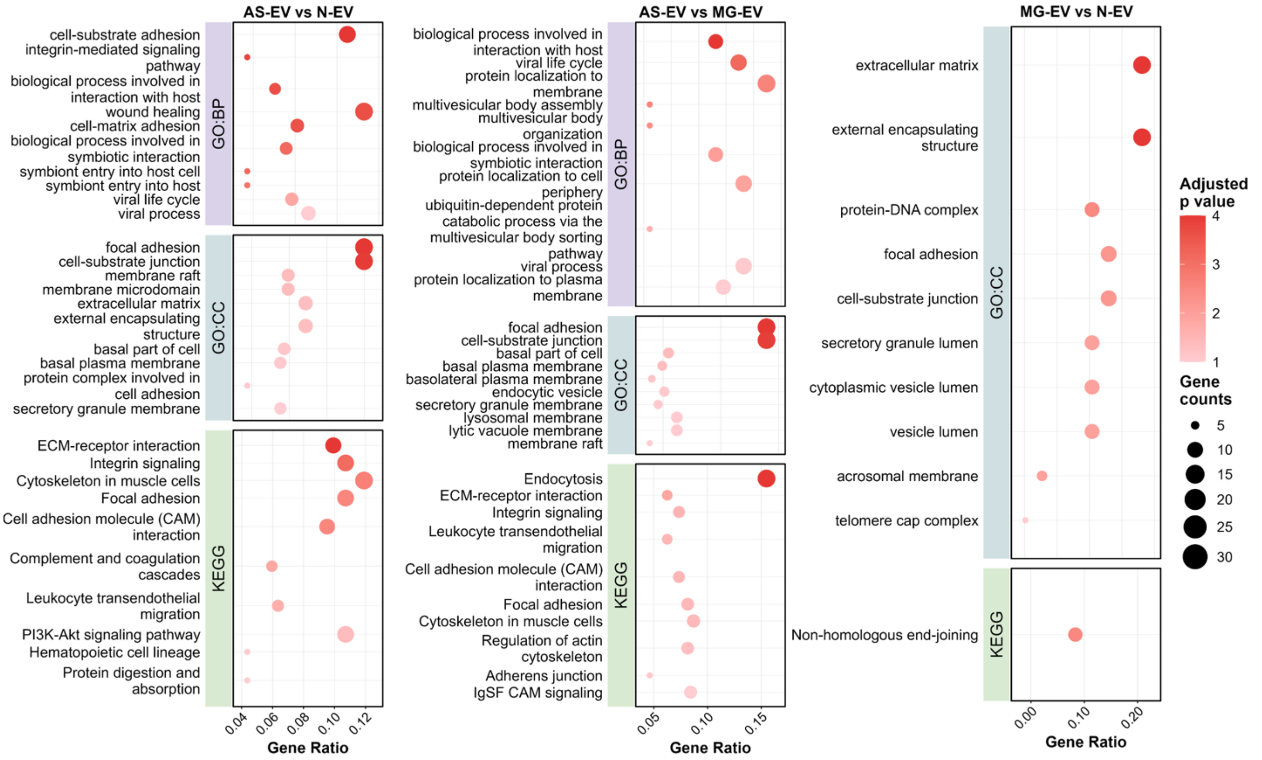


**Supplemental Figure** S2 GO:BP, GO:CC and KEGG pathway enrichment analysis on upregulated DEPs across pairwise EV comparisons. Dot plot showing up to top 10 GO:BP, GO:CC and KEGG pathway enrichment of upregulated DEPs in AS-EV vs N-EV, AS-EV vs MG-EV and MG-EV vs N-EV comparisons (adjusted p-value < 0.05). Dot size represents gene count and color indicates statistical significance ($-\log_{10}$(adjusted *p*-value)).

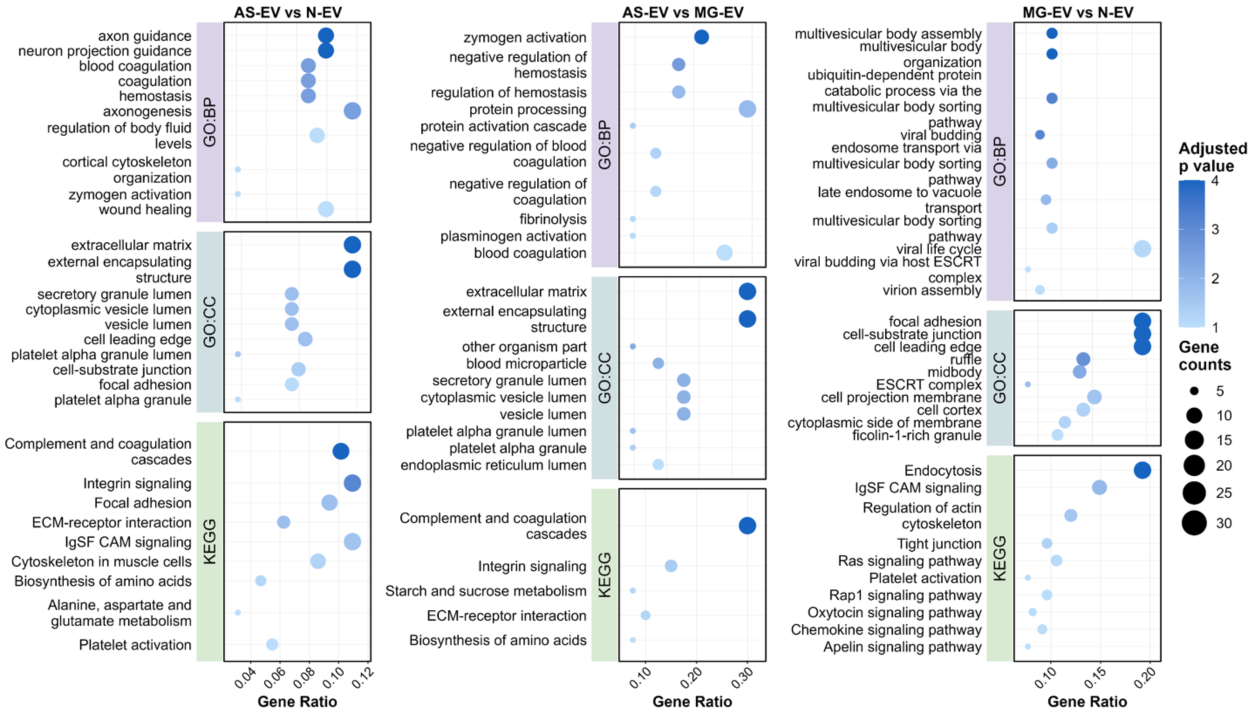


**Supplemental Figure S3** GO:BP, GO:CC and KEGG pathway enrichment analysis on downregulated DEPs across pairwise EV comparisons. Dot plot showing up to top 10 GO:BP, GO:CC and KEGG pathway enrichment of downregulated DEPs in AS-EV vs N-EV, AS-EV vs MG-EV and MG-EV vs N-EV comparisons (adjusted p-value < 0.05). Dot size represents gene count and color indicates statistical significance (-$\log_{10}$(adjusted *p*-value)).

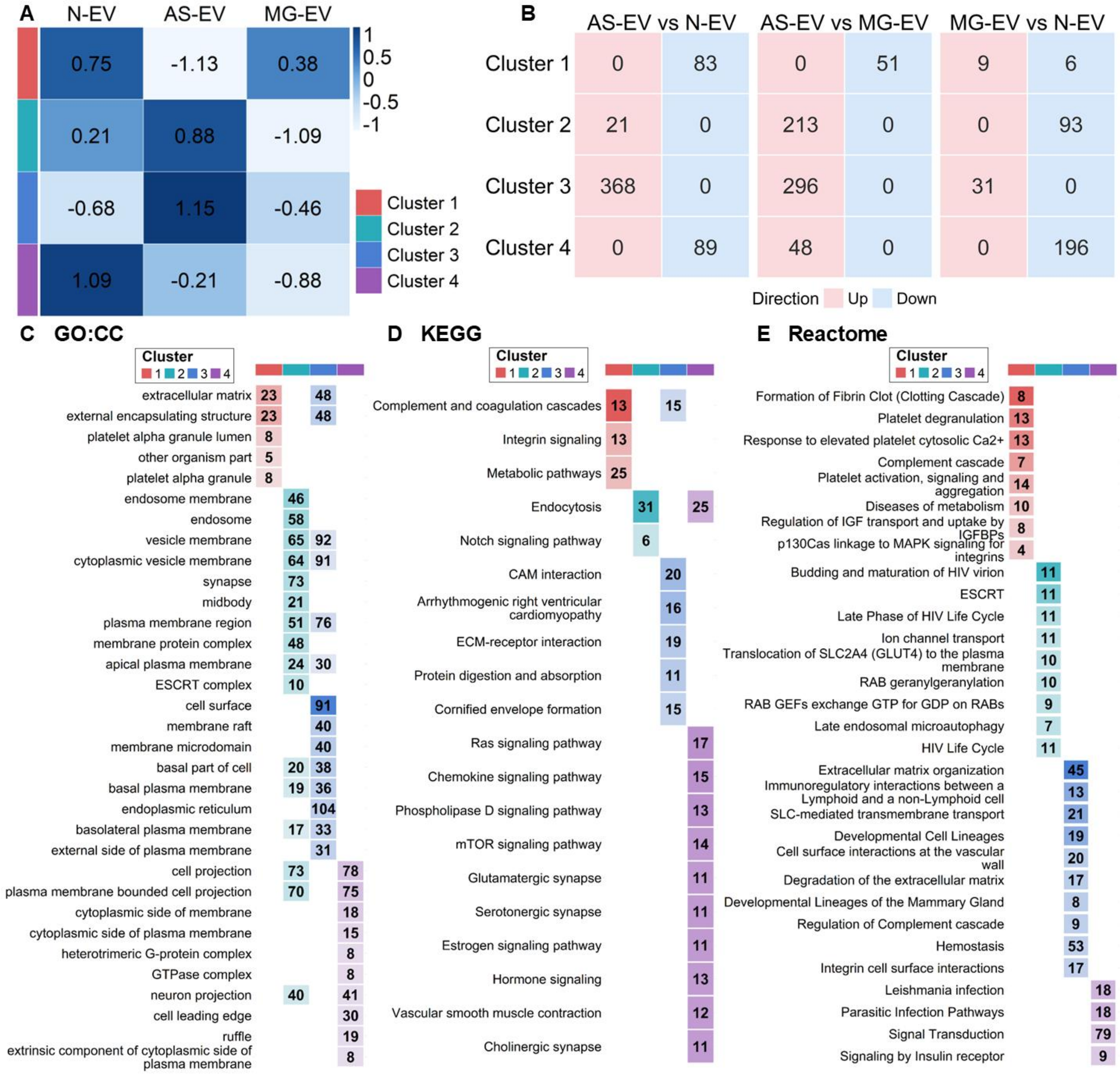


**Supplemental Figure S4** Protein abundance patterns, pairwise differential expression, and functional enrichment analysis of consensus clustering. **A.** Heatmap of row-scaled z-scores of mean protein abundance across N-EV, AS-EV, and MG-EV for each cluster. Values within cells indicate z-score magnitude. **B.** Tile plots showing the numbers of upregulated and downregulated DEPs in each cluster across three pairwise comparisons. Numbers within tiles indicate protein counts. **C-E.** Dot plots showing up to top 10 enriched (**C**) GO:CC terms, (**D**) KEGG pathways, and (**E**) Reactome pathways for each cluster (adjusted *p*-values< 0.05).

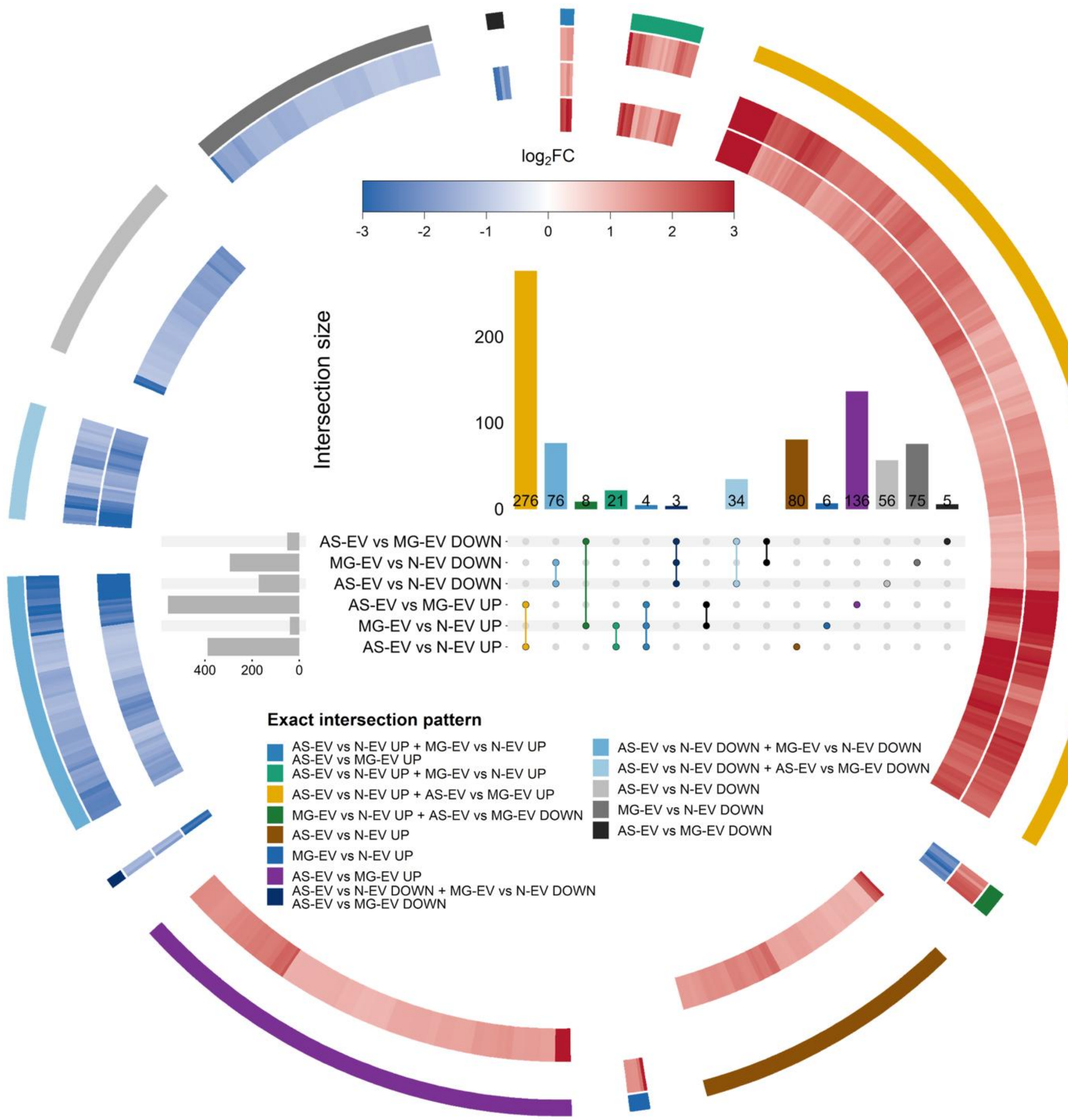


**Supplemental Figure S5** DEPs across pairwise comparisons. The UpSet plot showing direction-specific intersections of DEPs among the AS-EV vs N-EV, AS-EV vs MG-EV, and MG-EV vs N-EV pairwise comparisons. The accompanying circular heatmap displays protein abundance patterns within each intersection subset. From the inner to the outer rings, the heatmap represents AS-EV vs N-EV, AS-EV vs MG-EV, and MG-EV vs N-EV comparisons, respectively. Color intensity corresponds to $log_2FC$.
